# Biologically-informed Killer cell immunoglobulin-like receptor (KIR) gene annotation tool

**DOI:** 10.1101/2024.08.13.607835

**Authors:** Michael K.B. Ford, Ananth Hari, Qinghui Zhou, Ibrahim Numanagić, S. Cenk Sahinalp

## Abstract

Natural killer (NK) cells are essential components of the innate immune system, with their activity significantly regulated by Killer cell Immunoglobulin-like Receptors (KIRs). The diversity and structural complexity of KIR genes present significant challenges for accurate genotyping, essential for understanding NK cell functions and their implications in health and disease. Traditional genotyping methods struggle with the variable nature of KIR genes, leading to inaccuracies that can impede immunogenetic research. These challenges extend to high-quality phased assemblies, which have been recently popularized by the Human Pangenome Consortium. This paper introduces BAKIR (Biologically-informed Annotator for KIR locus), a tailored computational tool designed to overcome the challenges of KIR genotyping and annotation on high-quality, phased genome assemblies. BAKIR aims to enhance the accuracy of KIR gene annotations by structuring its annotation pipeline around identifying key functional mutations, thereby improving the identification and subsequent relevance of gene and allele calls. It uses a multi-stage mapping, alignment, and variant calling process to ensure high-precision gene and allele identification, while also maintaining high recall for sequences that are significantly mutated or truncated relative to the known allele database. BAKIR has been evaluated on a subset of the HPRC assemblies, where BAKIR was able to improve many of the associated annotations and call novel variants. BAKIR is freely available on GitHub, offering ease of access and use through multiple installation methods, including pip, conda, and singularity container, and is equipped with a user-friendly command-line interface, thereby promoting its adoption in the scientific community.

## 1 Introduction

Natural killer (NK) cells are critical components of the human innate immune system, playing a pivotal role in the body’s first line of defense against infections and tumors. A significant aspect of their regulatory mechanism involves the Killer cell Immunoglobulin-like Receptors (KIRs), which are expressed on the surface of NK cells and some T lymphocytes. The KIR gene complex, located on chromosome 19, spans approximately 150–200 kilobases and includes a diverse array of 17 genes, among which 15 are coding genes and 2 are pseudogenes. These genes exhibit considerable copy number variations among individuals, with haplotypes containing between 8 to 14 KIR genes which complicates their study but underscores their significance in genetic diversity and immune response [1–3].

KIRs interact with human leukocyte antigens (HLA) on the surfaces of target cells, influencing NK cell activation and inhibition. This dynamic balance between activating and inhibitory signals is crucial for NK cell function, affecting the development, tolerance, and activation of these cells [2]. The diversity of KIR haplotypes, classified into A (inhibitory) and B (activating) types, has been linked to various immunemediated diseases, the success of organ transplants, and the body’s response to infectious diseases like HIV and HCV [4–7]. Furthermore, KIR genes have been implicated in the susceptibility and progression of certain cancers, with research into anti-KIR monoclonal antibodies in conjunction with other immunotherapies showing promising results [8–10].

Accurately genotyping KIR genes from next-gen sequencing is a complex challenge exacerbated by the inherent diversity and structural variations present within the KIR locus. Even when utilizing high-quality, phased genome assemblies, researchers face significant obstacles. One of the primary issues is the heterogeneity of KIR haplotypes, which can vary dramatically between individuals, featuring different numbers and combinations of activating and inhibitory genes [11]. This variability complicates the genotyping process, as standard approaches based on fixed haplotype structures found in reference genomes may not accurately reflect the true diversity present within an individual’s genome [12]. Moreover, the presence of extensive copy number variations and structural variations within the KIR locus further complicates the analysis. Traditional methods such as GATK [13] that assume a diploid genome structure can lead to misinterpretations, overlooking or inaccurately representing these critical genomic features. These problems are shared with other immune-loci such as immunoglobulin heavy chain (IGH) and has led to the development of custom WGS genotyping approaches [14–16] and assembly annotation tools [17]. This highlights the necessity for genotyping tools that can adapt to the complexities of KIR genes, ensuring that annotations are reflective of the true genetic architecture.

The recent publication of the Human Pangenome Reference Consortium (HPRC) high-quality, haplotype-resolved genome assemblies [18] underscores the critical need for KIR genotyping annotations that transcend the limitations of fixed haplotype structures and conventional allele databases. These advanced genome assemblies offer an expansive view of human genetic variation, providing an ideal foundation for accurate KIR genotyping. However, to leverage this resource effectively, there is a need for genotyping tools capable of interpreting the nuanced genetic information contained within these assemblies. High-quality and dependable genotype annotations are essential for a deeper understanding of the KIR locus’s genetic variation. They play a crucial role in advancing immunogenetic research by providing a reliable benchmark for the development of both short- and long-read whole-genome sequencing-based genotyping tools. Accurate annotations serve as a critical resource for the scientific community, facilitating the validation and comparison of genotyping methodologies and enhancing our overall comprehension of KIR gene function and its impact on human health.

Addressing the complexities involved in KIR genotyping, we have developed BAKIR—Biologically-informed Annotator for KIR locus (pronounced /ba-ker/), a tool aimed at annotating KIR genotypes on high-quality, phased genome assemblies. BAKIR is structured to handle the complex genetic architecture of the KIR locus, allowing for the production of detailed genotype annotations. Such annotations are crucial for furthering our understanding of NK cell biology, along with its various implications for health and disease.

## 2 BAKIR

BAKIR takes as input a FASTA file containing a phased assembly sequence, either concatenated or split into contigs, and returns the start position, end position, closest allele, and variants of each KIR copy in the assembly. One of the key contributions of BAKIR is its approach to genotyping, which emphasizes the importance of “core variants”, which are variants that affect the downstream protein product or its expression. In most cases, the set of core variants is the set of non-synonymous variants, but it can also include other functionally-relevant variants such as those in the UTR. Unlike standard genotyping methods, which often rely on naive sequence mapping or simple alignment, BAKIR selects the best gene and allele by identifying and focusing on core variants. This approach ensures that the genotyping process is not only based on sequence similarity but also on biological and functional relevance, aligning closely with methods used, like those in pharmacogenomics, where functional variant-informed genotyping is widely accepted [19–21]. By comparing these core variants to those cataloged in the allele database, BAKIR can more accurately reflect the biological impact of different KIR alleles, providing a deeper and more meaningful understanding of individual and population-level genetic variability.

### 2.1 Methods

The BAKIR operates by initially fetching and analyzing the IPD-KIR allele database [22] to ensure comprehensive genomic representation of KIR alleles, including those only available in cDNA form. A key component of this analysis is determining the set of functionally-relevant core variants. It then implements a robust analytical framework to map these alleles to the provided haplotype-resolved genome assembly.

#### 2.1.1 Putative Gene Location

In its first stage, BAKIR aligns the database of KIR alleles with the genome assembly in question, utilizing `minimap2` [23]. Overlapped mappings are then consolidated to establish the preliminary gene locations in the assembly with the largest range. This approach allows for identification of gene copies that can harbor structural variations such as large deletions, as it integrates information from heavily clipped mappings, thereby ensuring a comprehensive identification of all possible KIR gene locations and setting the groundwork for subsequent analysis.

#### 2.1.2 Putative Gene Identification

Following the initial mapping, BAKIR’s next stage identifies the specific types of genes located at each putative gene site. This identification uses the prevalence of alleles mapped with each region, thus highlighting the most probable gene candidates for subsequent analysis. The wildtype sequence of the identified gene is then mapped back to the putative gene region of the assembly to refine the start and end locations of the gene in the assembly. This step ensures that the gene boundaries are accurately defined, preparing the groundwork for the subsequent variant calling stage.

#### 2.1.3 Variant Calling

In the variant calling stage, BAKIR compares the sequence of the wildtype allele of the matched gene to that of putative assembly genes. This comparison, performed through direct semi-global alignment with parasail [24], enables high-confidence variant calling. Variants are then classified into core and silent categories, with the former including missense, nonsense, and frameshift mutations, and the latter encompassing synonymous mutations and intronic variants. This is done by applying each variant to the wildtype sequence, translating the modified sequence, and determining if there is a change in the protein sequence.

#### 2.1.4 Putative Gene Location Refinement

At this stage the putative assembly gene location is refined, addressing any erroneous clipping that may have been generated by the mapping tool in the previous stages. In the case where prefix or suffix deletions (i.e., deletions that remove the head or tail of the gene sequence) have been called in the variant calling stage, the putative assembly gene sequence is re-extracted, taking into account these indel variants, and variants are re-generated by repeating the process defined in Section 2.1.3. If these new variants represent a less complex variant set than those originally generated, they are used in the subsequent steps, and the start and end positions of the putative gene in the assembly sequence are updated.

#### 2.1.5 Allele Calling

In the final stage, allele calling is performed. BAKIR selects the allele that most closely mirrors the identified gene sequence, based on calculating Hamming distance between the called core variants of the assembly and the core variants of each allele of the matched gene, and selecting the allele with the minimum distance. To calculate this distance, we first collect all core variants for all alleles of the given gene and compare the nucleotide values at these positions between a given candidate allele and the assembly gene copy. For easier interpretation we output the Hamming distance divided by the number of core variants for the gene, plus the number of novel core variants called for the assembly copy. Note that for indels, each position of the indel is considered independently.

In instances of identical scores, the tool applies a secondary distance measurement based on silent variants using the Jaccard distance. Utilizing this two-step similarity measurement approach ensures that the final annotation accurately reflects the true biological and functional characteristics of the gene, thereby facilitating precise and meaningful genotyping outcomes.

#### 2.1.6 Outputs

After genotyping, BAKIR provides two output files: a TSV file that provides a summary of the alleles called on a gene-by-gene basis, as well as an in-depth YAML file. This second file provides detailed comprehensive information about the allele called for every copy, as well as both matching and novel variants, and variant similarity metrics to the top 5 closest matching alleles.

## 3 Result

To evaluate BAKIR, we genotyped 52 phased HPRC assemblies from 26 samples and compared the annotations against those of *Immuannot* [25] and *SKIRT* [26]. BAKIR and SKIRT both identified 498 KIR copies across all assemblies, while Immuannot identified 497. A total of 28 / 498 and 31 / 497 copies differed in major (three digit) allele calls from those made by SKIRT and Immuannot respectively, resulting in allele call concordance of 94.2% and 93.9%. See Fig 1 for visualization of concordance on a per-gene basis. All of these differences in allele calls were checked manually to confirm the BAKIR calls by extracting the regions specified by BAKIR and the alternative tool, aligning them to wildtype, calling variants and performing a multiple-sequence alignment [27] of the extracted sequences and their called alleles (see Fig 2, Supplementary Data 1 for comparison of all discordant allele calls). In nearly all of the cases of discordant allele calls, SKIRT and Immuannot selected the allele with the lowest global alignment distance, while BAKIR chose the allele with more shared core variants and fewer novel core variants (see Table 1, Fig 2). In the remaining cases, the allele selected by BAKIR was both the closest in terms of core variant distance and global alignment distance.

**Table 1:** Example of differing allele calls made by BAKIR and Immuannot for a copy of KIR2DL4 on sample HG02630, maternal haplotype. While the Immuannot allele ∗005101 has a lower global edit distance, the assembly is missing a core variant that is also not present in the BAKIR choice of ∗01201 —see Fig 2 for a visualization and description of the alignment of these alleles.

| Tool | Allele call | Global edit distance | Assembly missing core variants | Assembly new core variants | Hamming distance core variants |
| --- | --- | --- | --- | --- | --- |
| BAKIR | KIR2DL4*01201 | 86 | None | None | 0.0 |
| Immuannot | KIR2DL4*0050101 | 23 | (10521, C>G) | None | 0.00813 |

**Figure 1:**
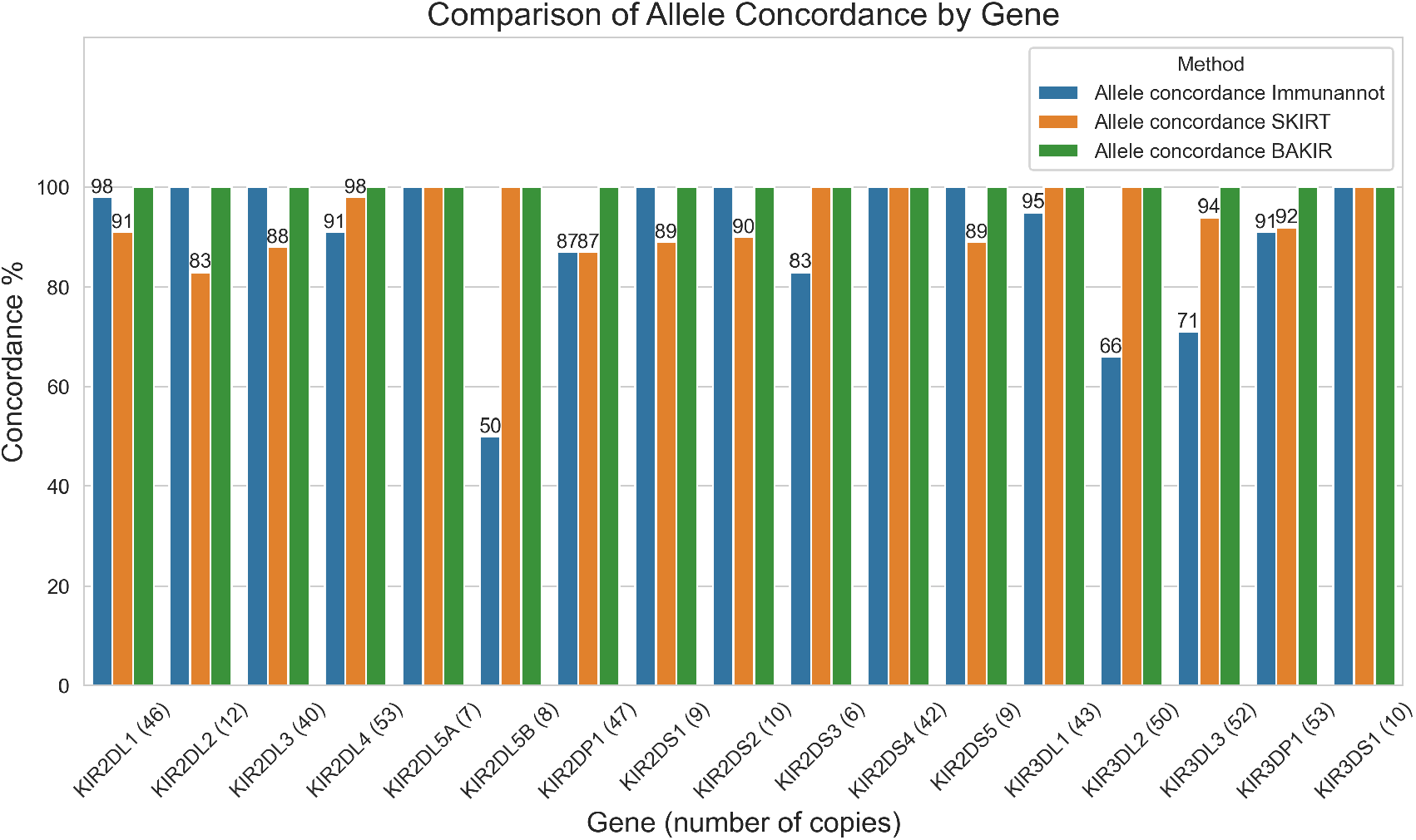
Summary of allele call concordance between BAKIR, Immuannot and SKIRT allele calls for the 52 HPRC assemblies for each KIR gene. Rather than depicting a comparison of absolute accuracy, this demonstrates the differences between the functionally-aware allele calls of BAKIR, based on core variants, and those of Immuannot and SKIRT. All BAKIR discordant alleles have been manually validated to be the best biologically-informed call (see Section 3).

**Figure 2:**
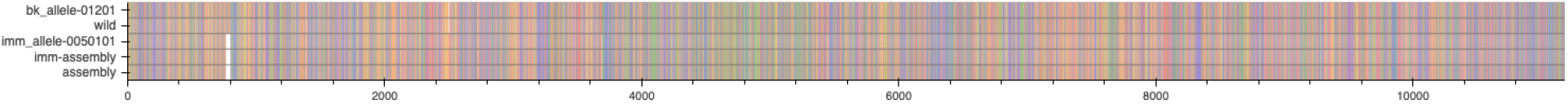
Alignment visualization for annotation described in Table 1. Note that the assembly sequence has an deletion shared by the ∗0050101 allele selected by Immuannot, near position 800, relative to the ∗02801 allele selected by BAKIR. However, this deletion, along with other variants contributing to the global edit distance, occur in introns and as a result are non-functional, while the allele defining C>G variant of ∗0050101 at wildtype position 10521 is not found in the assembly sequence, making ∗01201 a more functionally-relevant allele choice. `Wild` indicates the wildtype sequence, `assembly, imm-assembly` indicates the sequences from the assembly as found by BAKIR and Immuannot respectively (identical in this case), `bk_allele, imm_allele` indicate the sequences of the alleles chosen by BAKIR and Immuannot respectively.

## 4 Availability and Implementation

BAKIR can be found on GitHub (https://github.com/algo-cancer/bakir), and can be installed via pip, conda, docker, or singularity. It is run using a simple command-line interface, which can be queried by running bakir -h after installation. All code and scripts used to generate the results can be found at https://github.com/michael-ford/bakir-paper-methods.

## References

[1] Eric Vivier, Elena Tomasello, Myriam Baratin, Thierry Walzer, and Sophie Ugolini. Functions of natural killer cells. Nature immunology, 9(5):503–510, 2008.

[2] Katharine C Hsu, Shohei Chida, Daniel E Geraghty, and Bo Dupont. The killer cell immunoglobulin-like receptor (KIR) genomic region: gene-order, haplotypes and allelic polymorphism. Immunological reviews, 190(1):40–52, 2002.

[3] HG Shilling, K Lienert-Weidenbach, NM Valiante, M Uhrberg, and P Parham. Evidence for recombination as a mechanism for KIR diversification. Immunogenetics, 48:413–416, 1998.

[4] Salim I Khakoo and Mary Carrington. KIR and disease: a model system or system of models? Immunological reviews, 214(1):186–201, 2006.

[5] Jeanette E Boudreau, Tiernan J Mulrooney, Jean-Benoît Le Luduec, Edward Barker, and Katharine C Hsu. KIR3DL1 and HLA-B density and binding calibrate NK education and response to HIV. The Journal of Immunology, 196(8):3398–3410, 2016.

[6] Lies Boelen, Bisrat Debebe, Marcos Silveira, Arafa Salam, Julia Makinde, Chrissy H Roberts, Eddie CY Wang, John Frater, Jill Gilmour, Katie Twigger, et al. Inhibitory killer cell immunoglobulin-like receptors strengthen CD8+ T cell–mediated control of HIV-1, HCV, and HTLV-1. Science immunology, 3(29):eaao2892, 2018.

[7] Jing Li, Maxim Zaslavsky, Yapeng Su, Jing Guo, Michael J Sikora, Vincent van Unen, Asbjørn Christophersen, Shin-Heng Chiou, Liang Chen, Jiefu Li, et al. KIR+ CD8+ T cells suppress pathogenic T cells and are active in autoimmune diseases and COVID-19. Science, 376(6590):eabi9591, 2022.

[8] Holbrook E Kohrt, Ariane Thielens, Aurelien Marabelle, Idit Sagiv-Barfi, Caroline Sola, Fabien Chanuc, Nicolas Fuseri, Cécile Bonnafous, Debra Czerwinski, Amanda Rajapaksa, et al. Anti-KIR antibody enhancement of anti-lymphoma activity of natural killer cells as monotherapy and in combination with anti-CD20 antibodies. Blood, The Journal of the American Society of Hematology, 123(5):678–686, 2014.

[9] Francois Romagné, Pascale André, Pieter Spee, Stefan Zahn, Nicolas Anfossi, Laurent Gauthier, Marusca Capanni, Loredana Ruggeri, Don M Benson Jr, Bradley W Blaser, et al. Preclinical characterization of 1-7F9, a novel human anti–KIR receptor therapeutic antibody that augments natural killer–mediated killing of tumor cells. Blood, The Journal of the American Society of Hematology, 114(13):2667–2677, 2009.

[10] Laura Chiossone, Pierre-Yves Dumas, Margaux Vienne, and Eric Vivier. Natural killer cells and other innate lymphoid cells in cancer. Nature Reviews Immunology, 18(11):671–688, 2018.

[11] Markus Uhrberg, Nicholas M Valiante, Benny P Shum, Heather G Shilling, Kristin Lienert-Weidenbach, Brian Corliss, Dolly Tyan, Lewis L Lanier, and Peter Parham. Human diversity in killer cell inhibitory receptor genes. Immunity, 7(6):753–763, 1997.

[12] C T Watson and F. Breden. The immunoglobulin heavy chain locus: genetic variation, missing data, and implications for human disease. Genes & Immunity, 13(5):363–373, 2012.

[13] Geraldine A Van der Auwera and Brian D O’Connor. Genomics in the cloud: using Docker, GATK, and WDL in Terra. O’Reilly Media, 2020.

[14] Michael KB Ford, Ehsan Haghshenas, Corey T. Watson, and S. Cenk Sahinalp. Genotyping and copy number analysis of immunoglobulin heavy chain variable genes using long reads. iScience, 23(9):101508, 2020.

[15] Michael KB Ford et al. Immunotyper-sr: A computational approach for genotyping immunoglobulin heavy chain variable genes using short-read data. Cell systems, 13(10):808–816, 2022.

[16] Oscar L. Rodriguez et al. A novel framework for characterizing genomic haplotype diversity in the human immunoglobulin heavy chain locus. Frontiers in immunology, 11:2136, 2020.

[17] William D. Lees et al. Digger: directed annotation of immunoglobulin and t cell receptor v, d, and j gene sequences and assemblies. Bioinformatics, 40(3):btae144, 2024.

[18] Wen-Wei Liao, Mobin Asri, Jana Ebler, Daniel Doerr, Marina Haukness, Glenn Hickey, Shuangjia Lu, Julian K Lucas, Jean Monlong, Haley J Abel, et al. A draft human pangenome reference. Nature, 617(7960):312–324, 2023.

[19] Andrea Gaedigk, Magnus Ingelman-Sundberg, Neil A Miller, J Steven Leeder, Michelle Whirl-Carrillo, Teri E Klein, and PharmVar Steering Committee. The Pharmacogene Variation (PharmVar) Consortium: incorporation of the human cytochrome P450 (CYP) allele nomenclature database. Clinical Pharmacology & Therapeutics, 103(3):399–401, 2018.

[20] Ibrahim Numanagić, Salem Malikić, Michael Ford, Xiang Qin, Lorraine Toji, Milan Radovich, Todd C Skaar, Victoria M Pratt, Bonnie Berger, Steve Scherer, et al. Allelic decomposition and exact genotyping of highly polymorphic and structurally variant genes. Nature communications, 9(1):828, 2018.

[21] Ananth Hari, Qinghui Zhou, Nina Gonzaludo, John Harting, Stuart A Scott, Xiang Qin, Steve Scherer, S Cenk Sahinalp, and Ibrahim Numanagić. An efficient genotyper and star-allele caller for pharmacogenomics. Genome Research, 33(1):61–70, 2023.

[22] James Robinson, Jason A Halliwell, Hamish McWilliam, Rodrigo Lopez, and Steven GE Marsh. IPD—the immuno polymorphism database. Nucleic acids research, 41(D1):D1234–D1240, 2012.

[23] Heng Li. Minimap2: pairwise alignment for nucleotide sequences. Bioinformatics, 34(18):3094–3100, 2018.

[24] Jeff Daily. Parasail: SIMD C library for global, semi-global, and local pairwise sequence alignments. BMC bioinformatics, 17(1):1–11, 2016.

[25] Ying Zhou, Li Song, and Heng Li. Full resolution HLA and KIR genes annotation for human genome assemblies. bioRxiv, 2024.

[26] Tsung-Kai Hung, Wan-Chi Liu, Sheng-Kai Lai, Hui-Wen Chuang, Yi-Che Lee, Hong-Ye Lin, Chia-Lang Hsu, Chien-Yu Chen, Ya-Chien Yang, Jacob Shujui Hsu, and Pei-Lung Chen. Genetic diversity and structural complexity of the killer-cell immunoglobulin-like receptor gene complex: A comprehensive analysis using human pangenome assemblies. bioRxiv, 2023.

[27] J. D. Thompson, D. G. Higgins, and T. J. Gibson. CLUSTAL W: improving the sensitivity of progressive multiple sequence alignment through sequence weighting, position-specific gap penalties and weight matrix choice. Nucleic Acids Research, 22(22):4673–4680, 1994.

